# Legacy of warming and cover crops on the response of soil microbial function to repeated drying and rewetting cycles

**DOI:** 10.1101/2023.12.21.571204

**Authors:** Adetunji Alex Adekanmbi, Yiran Zou, Xin Shu, Giacomo Pietramellara, Shamina Imran Pathan, Lindsay Todman, Tom Sizmur

**Affiliations:** Department of Geography and Environmental Science, University of Reading, Whiteknights, PO Box 217, Reading, Berkshire RG6 6AH, UK; Department of Sustainable Land Management, University of Reading, UK; Department of Soil Science and Land Management, Federal University of Technology, Minna, Nigeria; Department of Agriculture, Food, Environment and Forestry, University of Florence, Firenze, 50144, Italy

**Keywords:** Soil respiration, Cover Crops, Climate Change, Warming, Drought, Stress, Perturbation, Adaptation, Resistance, Osmolytes

## Abstract

The response of soils to extreme weather events will become increasingly important in the future as more frequent and severe floods and droughts are expected to subject soils to drying and rewetting cycles as a result of climate change. These extreme events will be experienced against a backdrop of overall warming. However, farmers are adopting cover cropping as a sustainable management practice to increase soil organic matter, benefit soil health, and to increase the resilience of soils to help mitigate the impacts of climate change. We examined the legacy of warming and cover crops on the response of soil microbial function to repeated drying and rewetting cycles. We introduced open top chambers to warm the soil surface of a field plot experiment in which cover crops (single species monocultures and 4-species polycultures) were grown over the summer after harvest and before planting of autumn sown cash crops in a cereal rotation. Soil samples were collected from warmed and ambient areas of the experimental plots in spring, before harvesting the cereal crop. We quantified respiration (a measure of soil microbial function) with high-frequency CO_2_ flux measurements after 0, 1, 2, 4, or 8 wet/dry cycles imposed in the laboratory and the addition of barley grass powder substrate at a ratio of 10 mg g^-1^ soil. Cover crop mixtures created a negative legacy effect in the soil which resulted in lower cumulative substrate induced respiration than expected from the average of the same species grown in monoculture. Repeated drying and rewetting cycles increased the cumulative substrate induced respiration rate observed, suggesting that repeated perturbations selected for a community adapted to processing the barley shoot powder more quickly. This adaptation may have resulted in a greater osmolyte production or reacquisition by microorganisms exposed to repeated drought events. Osmolytes are rapidly metabolised upon re-wetting and may have primed the decomposition of the barley shoot powder to a greater extent in soils previously exposed to drying and rewetting cycles. When we calculated the cumulative respiration after 8 wet/dry cycles, relative to cumulative respiration after 0 wet/dry cycles (which we infer represents the extent to which microbial communities adapted to repeated drying and rewetting cycles) our data revealed that the legacy of warming significantly reduced, but cover crops significantly increased, soil microbial community adaptation. This adaptation of the soil microbial community was positively correlated with the concentration of water extractable organic carbon in the soils prior to imposing the drying and rewetting cycles and/or adding the substrate. The availability of labile carbon may have mediated the ability of microorganisms to synthesise osmolytes in response to drought. We conclude that cover crops may enhance the ability of the soil microbial community to adapt to drought events and mitigate the impact of warming, possibly due to the provision of labile organic carbon for the synthesis of osmolytes.

## Introduction

The Earth is warming at an increasing rate and warming of up to 6.4 °C is expected during the 21^st^ century if mitigation methods are not in place (Carey et al., 2018; Li et al., 2022). Warming could increase soil respiration, and increase CO_2_ flux from soils to the atmosphere, resulting in a positive feedback (Rustad et al., 2001; Bardgett et al., 2008; Dutta and Dutta, 2016). Field experiments which elevate soil temperatures by 0.3 to 6°C in various ecosystems, have consistently shown increases in soil respiration (Rustad et al., 2001; Kuffner et al., 2012). Winter warming has been observed to outpace summer warming in the northern hemisphere, and particularly in the UK (Vogelsang & Franses, 2005). Kreyling et al., (2019) showed that warming of soil by up to 1.7 °C from October to March affected several ecological processes, including plant performance, soil respiration and soil biological and chemical properties. A rise in temperatures of between 0.5 and 2.0 have also resulted in increases in soil pH and available phosphorus, but lower phosphatase, catalase, and urease activities (Guoju *et al*., 2012). Low-level winter warming was observed to improve the availability of soil carbon and nitrogen and increase the soil microbial biomass and greenhouse gas emissions (Liu *et al*., 2023) Understanding the response of the soil microbial community to warming may enhance our ability to predict the impact of climate change on soil respiration (Kreyling, 2010; Kreyling et al., 2019) and ecological processes more generally.

As a result of the overall increase in global temperatures, a greater frequency and severity of extreme weather events is being experienced globally (Rahmstorf & Coumou, 2011). In the UK, and much of Europe, this manifests as greater incidence of drought (Lavalle *et al*., 2009), causing concern about the resilience of soils and crop yields to perturbations caused by extreme weather events (Harkness *et al*., 2020). Therefore, our agroecosystems must be resilient to extreme events as well as higher temperatures. The ability of soil microbial communities to withstand and recover from perturbations and disturbances (including drought) is vital to ensure their continued ability to deliver multiple ecosystem services (Philippot *et al*., 2021). Long term strategies are required, in particular, to increase agricultural resilience to drought in temperate environments (Holman *et al*., 2021). When soil microorganisms are exposed to drought conditions, they increase their internal solute potential by synthesising low molecular weight compounds, osmolytes, which help to maintain homeostasis and turgor (Schimel, 2018). Upon re-wetting, a pulse of CO_2_ is often observed (Zhang *et al*., 2020), the origins of which have been much debated in the literature (Barnard *et al*., 2020). However, it is proposed that this flush may, in part, be due to the metabolism of osmolytes which primes the decomposition of other substrates (Warren, 2016). It is possible that repeated exposure of soil microbial communities to drought conditions may result in adaptation of the microbial community to better maintain soil function after drought (Allison, 2023), in much the same way that we observe thermal adaptation of soil microbial communities to warming (Bradford, 2013).

One important regenerative agriculture strategy which has recently surged in popularity in European cropping systems is to incorporate agro-ecological service crops, such as cover crops, into cropping rotations to enhance soil quality, encourage soil biodiversity, and reduce CO_2_ emissions (Papp et al., 2018; Radicetti et al., 2019). Such cover crops are planted as a subsidiary crop to a cash crop to enhance the overall conditions of the soil and it has been suggested that cover crops can help mitigate the impacts of climate change (Kaye & Quemada, 2017). Cover crops can enhance soil microbial biomass through carbon supplied by root exudates, shoots, and roots (Gyssels et al., 2005; Paterson et al., 2007; Calderón et al., 2016; Papp et al., 2018). Growing mixtures of cover crops, rather than single species monocultures, can result in greater increases in soil microbial functional diversity (Drost *et al*., 2020) and biomass (Shu *et al*., 2022) due to the provision of more diverse substrates. However, it is not known or understood whether cover crop mixtures (or monocultures) can increase the resistance and resilience of soil microbial communities to the effects of global warming, or extreme weather events.

Open Top Chambers (OTCs) are one of the most commonly used methods to invoke passive warming of soils in field experiments and have been demonstrated to effectively elevate both air and soil temperature in many terrestrial biomes (Carey et al., 2018; Sharkhuu et al., 2013). In this study we introduced OTCs over the 2019/20 and 2020/21 crop growing seasons (i.e. after cereal crops are planted until mid-spring) to warm the soil surface of an ongoing field plot experiment in which cover crops (single species monocultures and 4-species polycultures) were grown over the summer in between autumn sown cash crops in a cereal rotation. We collected soils samples in spring 2021 to compare soil microbial substrate-induced respiration in warmed and ambient (i.e. not warmed) plots where cover crops had been grown in monoculture, polyculture, or absent. We measured soil respiration using a high frequency respirometer after subjecting the soil to repeated (0, 1, 2, 4 or 8) drying and rewetting cycles and then applying a substrate of fresh barley shoot powder, following the assay described by Todman et al., (2018). We hypothesized that the legacy effect of warming would suppress substate-induced respiration after 8 repeated drying and rewetting cycles, relative to 0 cycles. We also hypothesised that cover crops would mitigate the impact of warming by supporting soil microbial activity, and that this effect would be more prevalent when cover crops were grown in polyculture than monoculture.

## Materials and Methods

### Field experiment design

The field plot experiment was located on the University of Reading farm at Sonning in Berkshire, UK and was established in August 2018. Each summer, after the harvest of a cereal crop, cover crops were planted and then terminated and incorporated prior to planting the next autumn sown cereal crop. The cereal rotation was Winter Wheat (2017/18); Winter Barley (2018/19); Winter Oats (2019/20); and Winter Wheat (2020/21). The cover crop treatments were control (no cover crops), buckwheat (*Fagopyrum esculentum*), berseem clover (*Trifolium alexandrinum*), oil radish (*Raphanus raphanistrum*), sunflower (*Helianthus annuus*), and a 4-species mixture of these four. The experiment was arranged in a randomised complete block design with 4 blocks. The field experiment included additional treatments not described here but this study focused on the 24 plots where cover crops were grown, and the residues incorporated into the soil. The selected plots are circled on the diagram shown in Figure S-1.

### Passive Warming

Open Top Chambers (OTCs) were installed after cereal crop establishment during the 2019/20 and 2020/21 growing seasons to act as passive warming devices by warming the soil under the OTCs. The OTCs were installed on 18^th^ December 2019 and 24^th^ November 2020, respectively, at crop seedling stage when about 5 leaves were unfolded. Each chamber was placed on the south-east end of each of the 24 plots to allow yield measurements to be made on the rest of the plot. Each OTC was a six-sided hexagon made from clear extruded Perspex acrylic plates with the following dimensions: 5 mm thickness with a 100 cm base, 57.74 cm top, 62 cm side cut at an angle 71.16° and each side was 50 cm high (Figure S-2). Our design for the OTC chamber conforms to the characteristics of the International Tundra Experiment (ITEX), with a similar shape and reinforcement as the hexagon chamber described by Marion et al., (1997). Clear Perspex acrylic sheets can transmit in excess of 92% of visible light and have higher light transmission capacity than glass. We monitored the effect of chambers on temperature within the top 10 cm of soil by inserting temperature probes coupled to Plus 2 Tinytag data loggers (Gemini, UK) in the centre of a chamber (warmed treatment) and at the equivalent location at the other end of the same plot (ambient treatment) of a control plot. The loggers were set to log temperature every 15 minutes while the OTCs were in place. The OTCs warmed the soil by, on average, 0.75±0.92 °C in the control plot in 2020/21. The difference in soil temperature under the OTC, compared to the ambient measurements is shown in Figure S-4.

### Soil sampling

We sampled soils on 17^th^ May 2021 from the top 10cm underneath the OTSs (warmed treatment), and the equivalent location at the other end (ambient treatment) of each plot circled in Figure S-1 using a trowel. The soil samples were sieved, fresh, to 4mm and divided into four subsamples. The first subsample was refrigerated at 4°C prior to KCl extraction to determine NH_4_^+^ and NO_3_^-^ availability. The second subsample was used for DNA extraction, and quantification of gene abundance of microbial communities using real time qPCR. The third subsample was air-dried for soil chemical analysis. The fourth subsample was used to assess the response of soil microbial respiration following repeated drying and rewetting cycles and addition of barley shoot powder as a substrate.

### Soil chemical analysis

Soil pH was determined by shaking soil samples with deionised water (1:2.5 mass/volume ratio) for 10 min and leaving the mixture to stand for 2 min before pH was measured using a digital type DMP-2 mV/pH meter (Thermo Orion). Total N and C concentrations were determined using C/N Elemental Analyser (Thermo Flash 2000 EA). The C/N ratio was then calculated from total C and N. NH_4_^+^ and NO_3_^-^ was extracted in 1 M KCl and then analysed using a Continuous Flow Analyzer (San++ Automated Wet Chemistry Analyzer - SKALAR). Available N was calculated from the sum of extractable NH_4_^+^ and NO_3_^-^. Moisture content and loss on ignition were determined by weight loss at 105 °C and 500 °C, respectively. Hot water extractable carbon (HWEOC) and cold water extractable carbon (CWEOC) were analysed as described by Ghani et al., (2003). Approximately 3 g air-dried soil samples of known moisture content were accurately weighed into 50 ml polypropylene centrifuge tubes. Thirty ml of ultra-pure water was then added to each tube before mixing on a rotary shaker at 30 rpm for 30 min at 20℃. This was followed by centrifuging at 3500 rpm for 20 min at 20 °C. The supernatants were then removed using polypropylene syringes and passed through 0.2 µm cellulose nitrate membrane filters into polypropylene universal tubes, discarding the first 3 ml of the filtrate each time. A further 30 ml of ultra-pure water was then added to each centrifuge tube before vortexing for 10 seconds and leaving in an 80 ℃ water bath overnight and the supernatants removed, as described above. Both supernatants were analysed for CWEOC and HWEOC, respectively using a Shimazu TOC analyser.

### DNA extraction and qPCR

Total soil DNA was extracted from 230 mg of soil using the DNeasy® PowerSoil® kit (Qiagen). Genomic DNA concentrations were estimated by a NanoDrop 2000 spectrophotometer (Thermo Fisher Scientific, USA) and yield of soil-extracted DNA was checked by 1 % agarose gel electrophoresis and quality control. The abundance of total bacterial 16S rRNA and total fungal ITS genes as well as bacterial and fungal beta-glucosidases genes were quantified using real-time PCR technique. Specific primers were used to amplify each targeted gene sequences. Primer sets and references reporting the cycling conditions are shown in Table S-1. In addition, the melting curve conditions were performed from 55 to 95 °C, with increments of 0.5 °C every five seconds. Each 25 µl of PCR reaction contained 40 ng of the DNA, 400 nM of each primer, and 12.5 µl 2x ITAQ SYBER Green Super mix (Bio-Rad, Munich, Germany). Standard curves were constructed using a purified PCR product of known concertation from pure culture, *Bacillus subtilus* and *Trcichoderma reesei* for eubacteria (16S rRNA genes) and fungi (ITS genes) respectively. Standard curves for fungal and bacterial beta-glucosidases for GH1 (glycoside hydrolase family 1) and GH3 (glycoside hydrolase family 3) were constructed using a purified PCR products of known concentration of common DNA mixture (equal amount of DNA from all samples collected in this study). The concentration of each purified product was measured by Picodrop Microliter UV/Vis Spectrophotometer. Ten-fold dilutions ranging from 10^0^ to 10^-7^ ng per lµL^-1^ were applied for the standard curve construction. All standards and samples were run in triplicate and duplicate, respectively. Quantification was performed on a CFX ConnectTM Real-Time PCR detection System using CFX ManagerTM software 3.1 (Bio-Rad, Munich, Germany). Further, quantified DNA concentration (ng/g soil) of each sample was converted into units of copy number per g soil according to SC method (Brankatschk *et al*., 2012).

### Imposition of drying and rewetting cycles and substrate induced respiration

The water holding capacity of each soil sample was determined and soils were adjusted to 85% of their water holding capacity and pre-incubated at approximately 22°C for one week prior to the drying and rewetting assay to avoid artefacts caused by the soil preparation. The assay is based on the approach described by Todman et al., (2018) and Fraser et al., (2016). To undertake the assay, the equivalent of 10 g (dry weight) of six fresh subsamples of each soil sample were weighed into tin trays (prior to pre-incubation). To one subsample (0 wet/dry cycles), 100 mg of barley shoot powder (C/N ratio 23.3) was added and mixed following (Kuan *et al*., 2007). After mixing, CO_2_ evolution was measured continuously for 96 hours at hourly time intervals using an automated multichannel respirometer and an EGA60 multi-sample gas exchange system (ADC Bioscientific Ltd). Samples that had not received substrate were exposed to 1, 2, 4, or 8 drying and rewetting cycles prior to the addition of barley shoot powder and substrate induced respiration being measured. Each drying and rewetting cycle consisted of 3 days of drying by enclosing samples in a sealed chamber with silica gel desiccant, followed by weighing and rewetting to 85% of the water holding capacity where they were maintained for a further 4 days, following Todman et al., (2018), as depicted in Figure S-5. It is important to note that the barley shoot powder was only added once to each sub-sample, so the assay measured the CO_2_ flux upon addition of a novel substrate after being exposed to 0, 1, 2, 4, or 8 wet/dry cycles (Figure 1). CO_2_ respired in (ppm) was converted to C-CO_2_ (µg g^-1^ soil h^-1^) and the cumulative sum (µg g^-1^ soil) of C-CO_2_ respired per sample over the 96 hours was calculated. We then quantified cumulative respiration after 8 wet/dry cycles as a percentage of the cumulative respiration of a sub-sample from the same plot after 0 wet/dry cycles to represent the extent to which the microbial community adapted to the drying and rewetting cycles.

**Figure 1.**
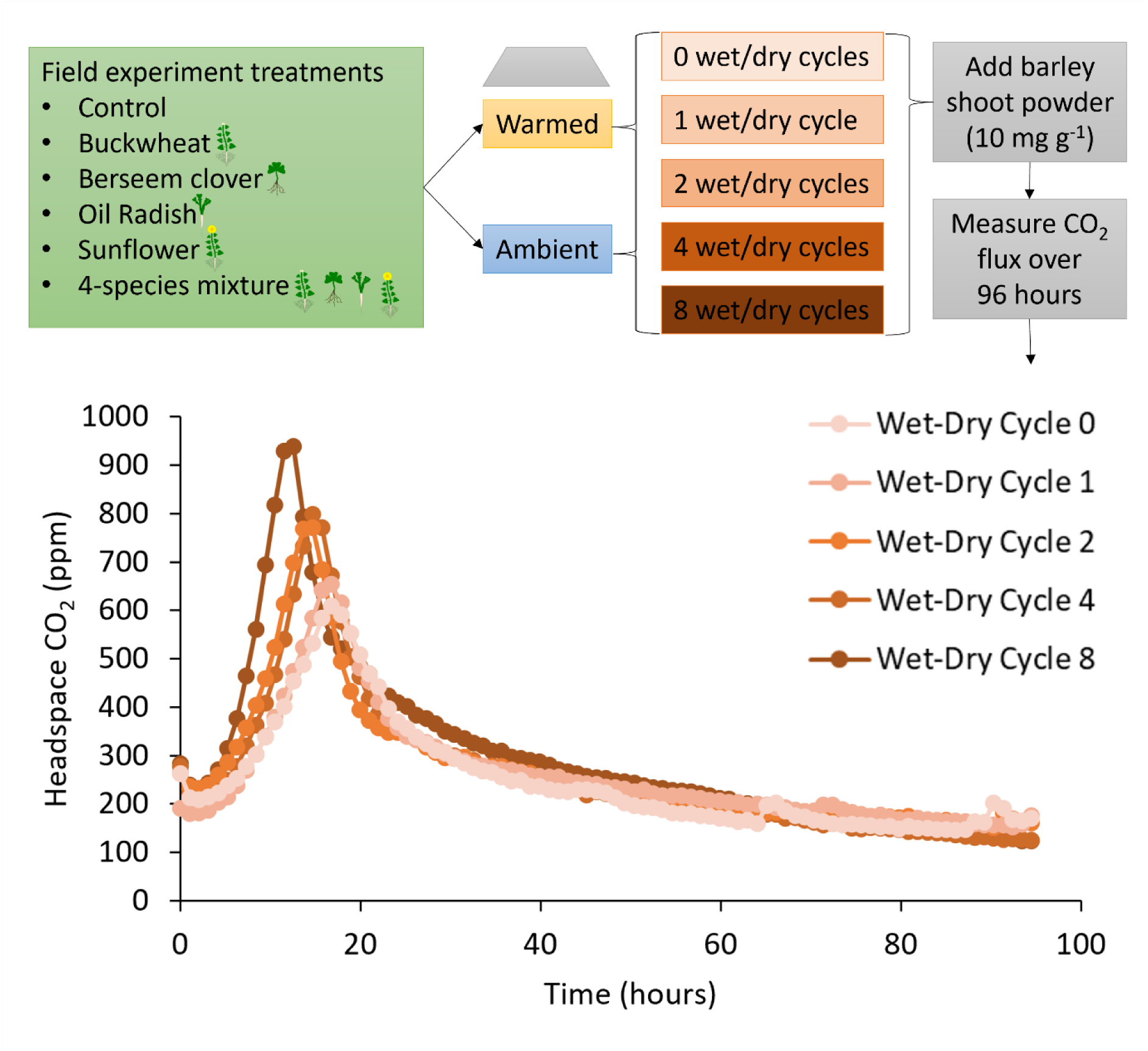
Schematic of the experimental design and procedure and average CO_2_ concentrations measured during the 96 hours after barley shoot powder addition plotted as an average of all 48 experimental treatments.

### Statistical Analysis

The effect of experimental treatments (passive warming and cover crops) on cumulative respiration (and the percentage of cumulative respiration after 8 wet/dry cycles relative to 0 wet/dry cycles) was analysed using the following ANOVA model: Warming*Cover crop/(Mix/Type) in GenStat (2021 version). The *Warming* factor had two levels (warmed and ambient). The *Cover crop* factor also had two levels (control and cover crops). The *Mix* factor had two levels (monoculture and 4-species mixture) with a dummy level representing the control treatment. The *Type* factor had four levels (buckwheat, berseem clover, oil radish, and sunflower) with dummy levels representing the control and mixture treatments. The relationships between the percentage of cumulative respiration after 8 wet/dry cycles relative to 0 wet/dry cycles and soil chemical and biological properties were analysed using Pearson’s correlation. The extent to which soil chemical and biological properties predict the cumulative respiration after 8 wet/dry cycles relative to 0 wet/dry cycles was explored using stepwise multivariate regression (stepwise selection of terms was based on p < 0.15).

## Results

### Differences in soil properties due to warming and cover crops

We observed several differences in the soil chemical and biological properties that can be attributed to the passive warming or cover crop treatments. There were no significant effects of *Warming*, *Cover Crop*, *Mix*, or *Type* on HWEOC, CWEOC, %LOI, extractable NO_3_^-^, C/N ratio or abundance of total bacteria (16S rRNA gene), total fungi (ITS region gene) or genes encoding a bacterial β-glucosidase enzyme (GH1) in soils (Table 1). However, the presence of cover crops significantly (*Cover crop*: P < 0.05) decreased soil pH (Figure S-6), %C (Figure S-7), and the abundance of genes encoding fungal β-glucosidase enzymes (GH3) (Figure S-11). There were no significant differences in soil properties between plots with the cover crop mixture and the average of the four monoculture plots (*Mix;* Table 1). However, cover crop species had a significant (*Type*: P < 0.05) effect on %N and the abundance of genes encoding both fungal and bacterial β-Glucosidase enzyme (GH3). Soils from plots planted with clover had the greatest %N (Figure S-8) while radish plots had the highest abundance of the bacterial GH3 gene (Figure S-9) and the lowest abundance of the fungal GH3 gene (Figure S-11). Warming significantly (*Warming*; P < 0.05) increased the abundance of genes encoding Fungal β-Glucosidase (GH1) (Table 1; Figure S-10).

**Table 1:** Summary table showing F values for nested ANOVA testing the effect of statistical factors (*Warming*, *Cover crops*, *Mix*, and *Type*) on soil properties.

|  | Warming | Cover<br>crops | Mix | Type |
| --- | --- | --- | --- | --- |
| HWEOC ($\mu\text{g g}^{-1}$) | 1.50 | 0.10 | 1.45 | 0.67 |
| CWEOC ($\mu\text{g g}^{-1}$) | 0.99 | 0.15 | 0.58 | 2.15 |
| Soil pH | 0.02 | 4.39* | 0.00 | 1.25 |
| % LOI | 0.06 | 0.87 | 0.01 | 0.71 |
| Extractable $\text{NO}_3^-$ (mg/kg) | 0.45 | 0.30 | 0.23 | 0.39 |
| Total Extractable N (mg/kg) | 0.68 | 0.06 | 0.14 | 0.22 |
| %N | 0.13 | 0.07 | 0.02 | 3.37* |
| %C | 2.45 | 4.38* | 0.13 | 1.17 |
| C/N | 0.60 | 1.25 | 0.15 | 2.14 |
| Bacteria abundance (16S rRNA gene<br>copies $\text{g}^{-1}$) | 0.11 | 4.29 | 1.27 | 2.30 |
| Fungal abundance (ITS gene copies<br>$\text{g}^{-1}$) | 0.28 | 2.42 | 0.59 | 0.31 |
| Bacterial β-glucosidase (GH1 gene<br>copies $\text{g}^{-1}$) | 2.49 | 4.24 | 0.41 | 1.06 |
| Bacterial β-glucosidase (GH3 gene<br>copies $\text{g}^{-1}$) | 3.98 | 0.01 | 0.31 | 4.52* |
| Fungal β-glucosidase (GH1 gene<br>copies $\text{g}^{-1}$) | 5.03* | 0.14 | 0.78 | 0.25 |
| Fungal β-glucosidase (GH3 gene<br>copies $\text{g}^{-1}$) | 1.10 | 13.91*** | 1.21 | 3.68* |

\* = $p < 0.05$ ; \*\* = $p < 0.01$ ; \*\*\* = $p < 0.001$

### Substrate induced respiration

Cumulative substrate induced respiration was significantly (*Mix*: P < 0.05) lower in the soils planted with a quaternary mixture of cover crop species than the average of the four monoculture plots after all five drying and rewetting cycle treatments (0, 1, 2, 4, and 8 cycles), as shown in Figure 2. The cover crop species planted also had a significant (*Type*: P < 0.05) effect on the substrate induced respiration, regardless of wet/dry cycle treatment (Figure 2). Generally, oil radish and sunflower plots resulted in greater substrate induced microbial respiration than buckwheat and clover plots. The effect of the *Mix* factor was always greater than the effect of the *Type* factor on cumulative substrate induced respiration. When data from each drying and rewetting cycle was statistically analysed on its own, there was no significant effect (P > 0.05) of *Warming* or *Cover crops* on cumulative substrate induced respiration.

**Figure 2:**
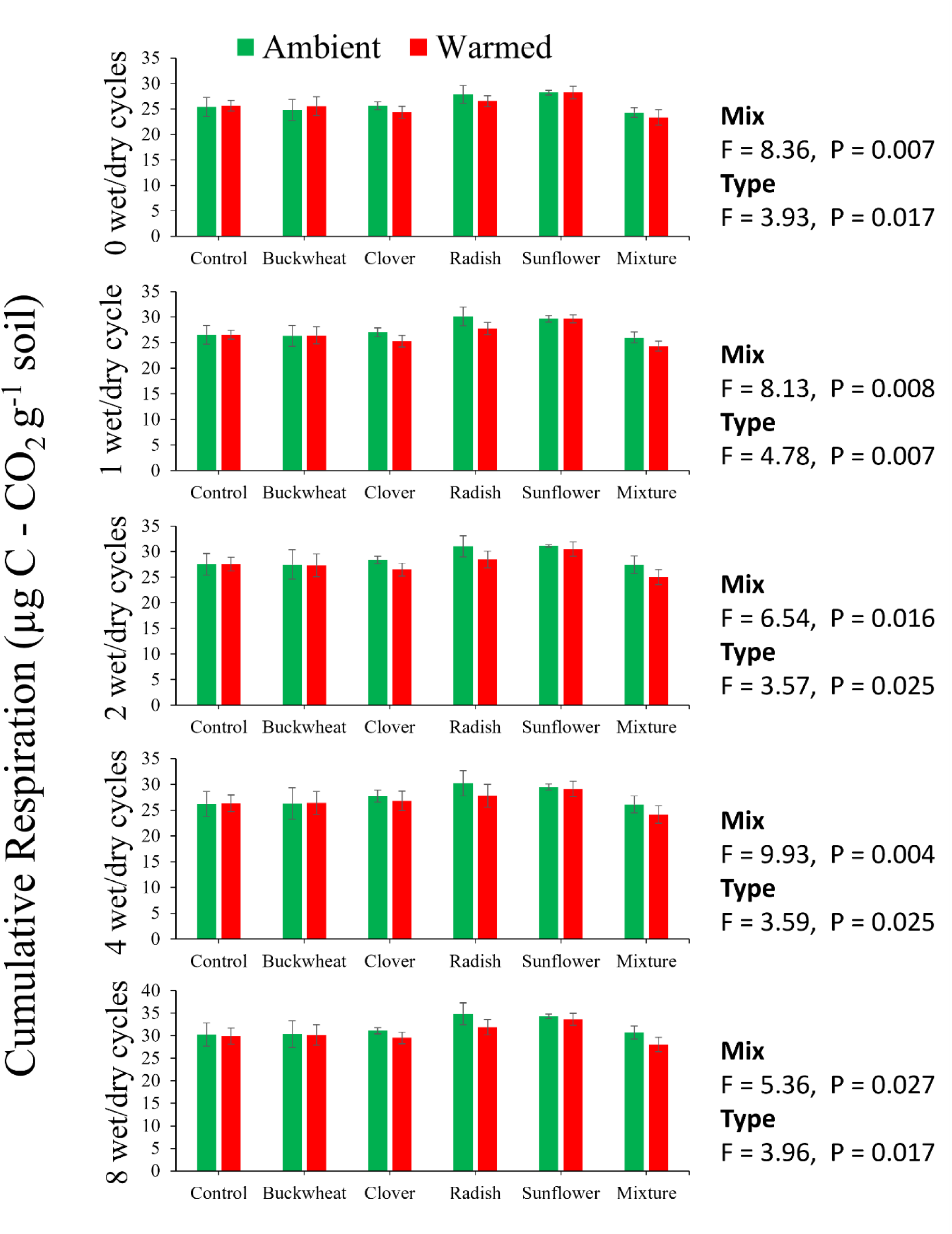
Cumulative substrate-induced microbial respiration after 0, 1, 2, 4 or 8 wet dry cycles from soils subjected to warmed or ambient treatments and planted with no cover crops (control), cover crop monocultures (buckwheat, clover, radish, and sunflower), or a 4-species cover crop mixture. *Warming* and *Cover crop* had no significant effect on cumulative respiration, but cover crop diversity (*Mix*) and species (*Type*) had a significant effect on soils subjected to each cycle. Error bars are standard deviations; n = 4.

In soils taken from all plots, more dry/wet cycles resulted in an earlier peak of CO_2_ flux, a higher maximum respiration rate, and greater cumulative substrate induced respiration, as demonstrated in Figure 1. Hence, a higher cumulative respiration was observed after 8 wet/dry cycles, relative to 0 wet/dry cycles, in contrast with our hypothesis. The largest impacts of drying and rewetting on substrate induced respiration were observed within the first 20 hours after the substrate was added to the soil. This observation is similar to that of Fraser *et al*., (2016a) who observed that the majority of the rapidly released CO_2_ from substrate-induced respiration following drying and rewetting occurred within the first 24 hours. Most of the C mineralised after substrate addition has been attributed to free sugars, soluble organic-N and fructans, and plant residues containing more of these compounds will lead to greater respiration compared to those with less concentrations (Gunnarsson *et al*., 2008; Fraser *et al*., 2016b). Warming significantly (*Warming*: P < 0.05) decreased the extent to which the cumulative respiration increased after 8 wet/dry cycles relative to 0 wet/dry cycles (Figure 3). However, Cover crops significantly (*Cover crop*: P < 0.05) increased the extent to which the cumulative respiration increased after 8 repeated dry/wet cycles relative to 0 dry/wet cycles (Figure 3). While the 4-species mixture of cover crops resulted in a slightly greater increase in cumulative respiration after 8 cycles, relative to 0 cycles, this was not statistically significant (*Mix*: P > 0.05) and there was no statistically significant difference (*Type*: P > 0,05) observed due to the four cover crop monoculture treatments (Figure 3).

**Figure 3:**
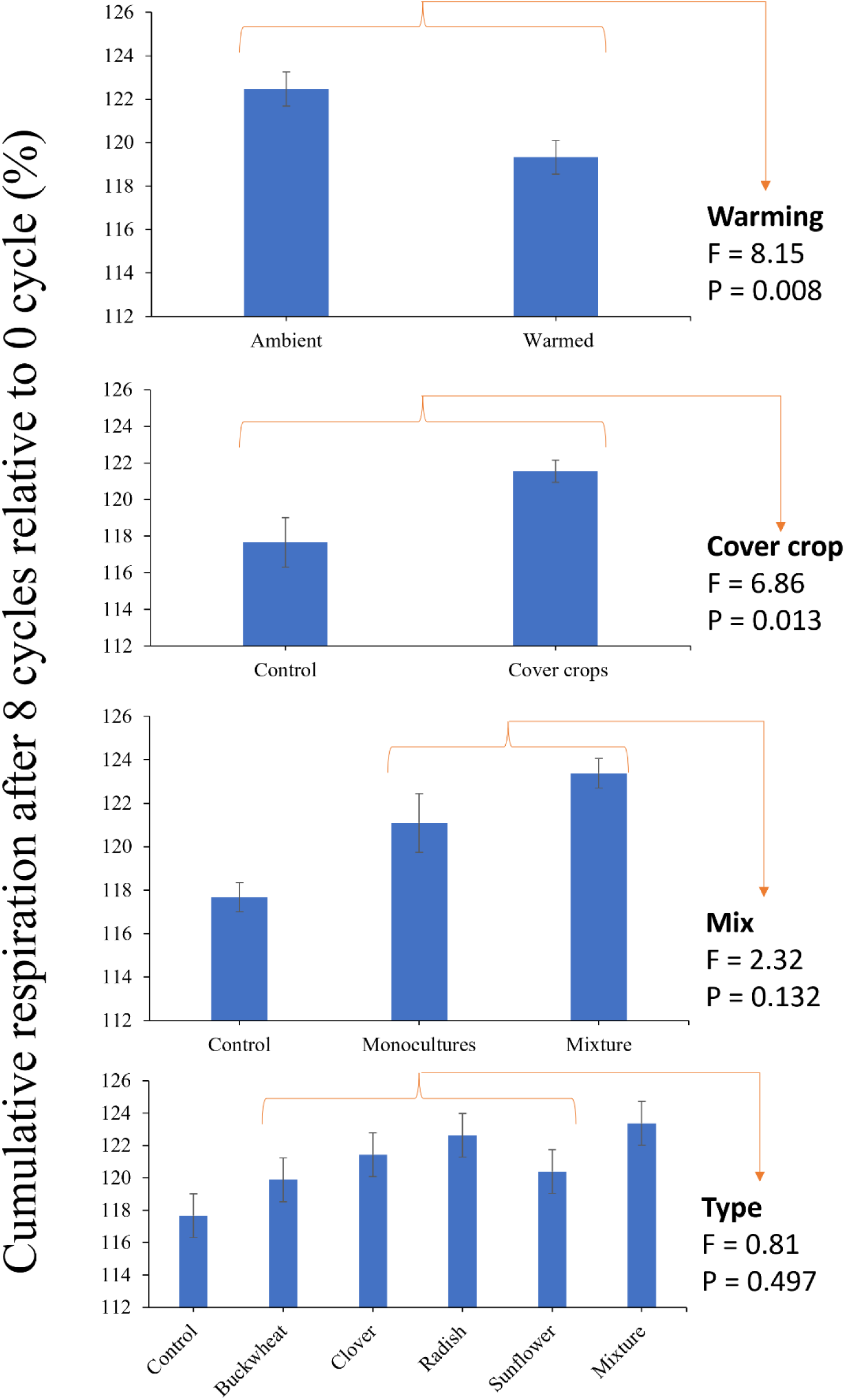
Effects of statistical factors (*Warming*, *Cover crops*, *Mix*, and *Type*) on cumulative substrate induced microbial respiration after 8 wet/dry cycles expressed as a percentage of the cumulative substrate induced microbial respiration observed after 0 wet/dry cycles. Error bars are standard deviations; n= 4.

### Relationship between cumulative substrate-induced respiration and soil properties

Correlation coefficients for the relationships between soil chemical and biological properties and the cumulative respiration after 8 wet/dry cycles relative to cumulative respiration after 0 wet/dry cycles are presented in Table 2. The relative increase in microbial respiration was significantly positively correlated (r = 0.493; P < 0.001) with cold water extractable organic carbon, and significantly negatively correlated (r = −0.414; P = 0.004) with the abundance of the Bacterial-GH1 gene encoding beta glucosidase and also negatively correlated, albeit not statistically significantly (r = −0.280; P = 0.057), with the abundance of the Bacterial-GH3 genes encoding beta glucosidase (Table 2). Stepwise multivariate regression revealed only one soil chemical or biological property (cold water extractable organic carbon) that significantly predicted (R^2^ adj. = 36.82%; P < 0.001) the relative increase in microbial respiration after 8 wet/dry cycles relative to 0 wet dry cycles, with no other property significantly improving the regression model (Figure 4).

**Figure 4.**
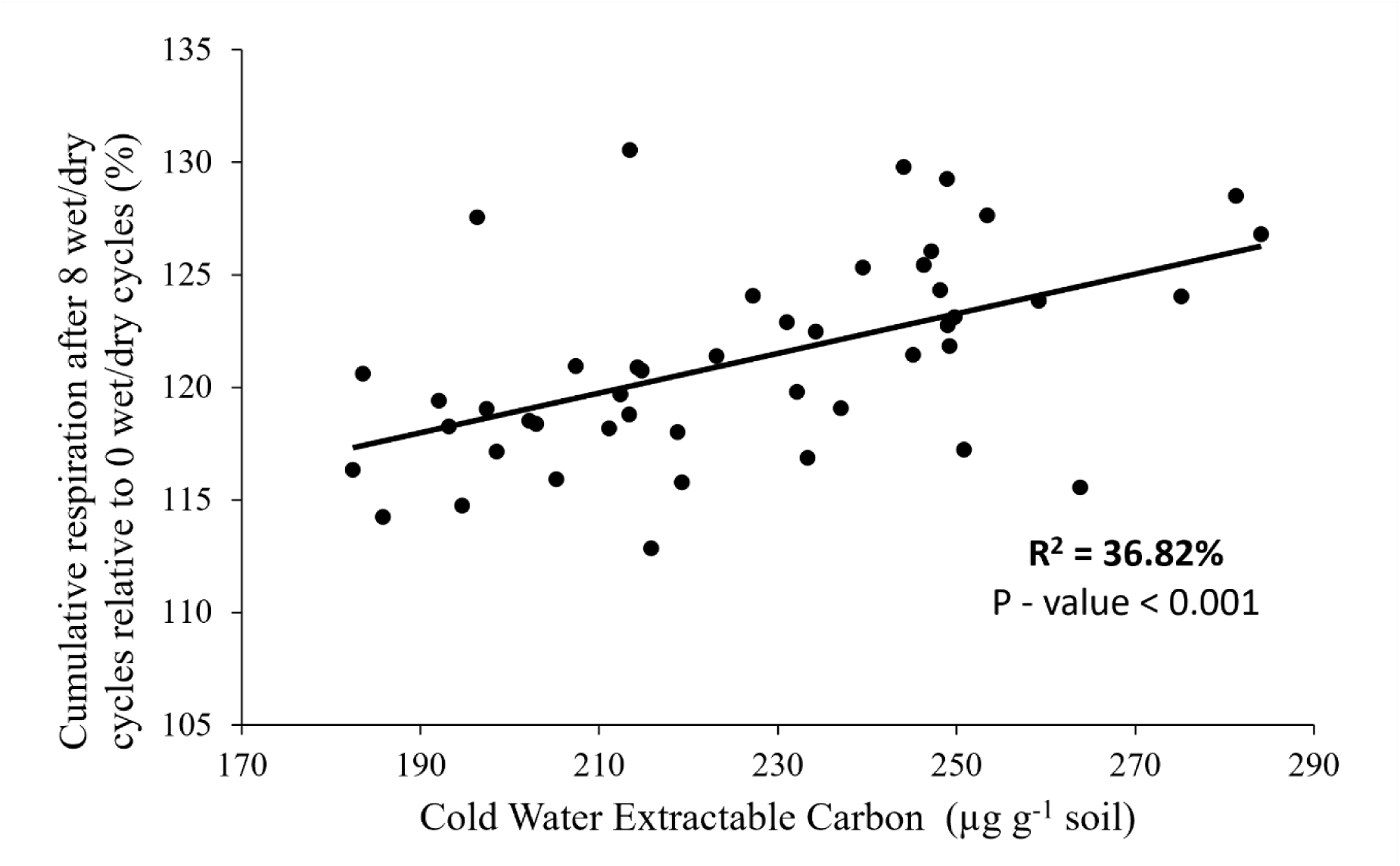
Regression between the cumulative respiration after 8 wet/dry cycles relative to cumulative respiration after 0 wet/dry cycles and cold water extractable organic carbon (CWEOC) for all samples and experimental treatments combined (n = 48).

**Table 2:** Pearson correlation coefficients and p values for the relationships between the cumulative respiration after 8 wet/dry cycles relative to cumulative respiration after 0 wet/dry cycles and soil chemical and biological parameters.

| Soil properties | Coefficient (r) | P value |
| --- | --- | --- |
| HWEOC<sup>a</sup> ($\mu\text{g g}^{-1}$) | 0.041 | 0.784 |
| CWEOC<sup>b</sup> ($\mu\text{g g}^{-1}$) | 0.493 | <0.001 |
| Soil pH | -0.040 | 0.789 |
| % LOI<sup>c</sup> | -0.103 | 0.492 |
| Extractable NO<sub>3</sub><sup>-</sup> (mg/kg) | -0.196 | 0.187 |
| Total Extractable N (mg/kg) | -0.206 | 0.164 |
| %N | -0.113 | 0.450 |
| %C | -0.168 | 0.260 |
| C/N | -0.048 | 0.747 |
| Bacteria abundance (16S rRNA gene copies g<sup>-1</sup>) | -0.165 | 0.268 |
| Fungal abundance (ITS gene copies g<sup>-1</sup>) | -0.260 | 0.077 |
| Bacterial β-glucosidase (GH1 gene copies g<sup>-1</sup>) | -0.414 | 0.004 |
| Bacterial β-glucosidase (GH3 gene copies g<sup>-1</sup>) | -0.280 | 0.057 |
| Fungal β-glucosidase (GH1 gene copies g<sup>-1</sup>) | 0.007 | 0.960 |
| Fungal β-glucosidase (GH3 gene copies g<sup>-1</sup>) | -0.277 | 0.059 |

<sup>a</sup> Hot Water Extractable Carbon; <sup>b</sup> Cold Water Extractable Carbon; <sup>c</sup> Loss on Ignition

## Discussion

### The influence of cover crop species diversity on substrate induced respiration

Our finding that the average legacy effect of planting 4 monocultures of cover crops resulted in significantly greater cumulative substrate induced respiration after all five drying and rewetting cycles treatments (Figure 2) implies that increasing plant diversity decreases soil microbial activity, in contrast to previous observations (Shu *et al*., 2021). Where positive relationships between plant diversity on ecosystem function are observed in field experiments, they can often be explained by greater plant biomass in more diverse plant communities (Zak *et al*., 2003). None of the soil chemical or biological parameters measured were significantly affected by cover crop diversity (apart from substrate induced respiration) to aid our mechanistic explanation of this result.

### The influence of cover crop species type on substrate induced respiration

Rhizodeposition during the growth phase of the cover crop and residue decomposition after cover crop termination and incorporation both regulate cover crop legacy effects on the soil microbiome (Spedding et al., 2004; Nannipieri et al., 2023). Cover crop-induced legacy effects on the soil microbiome can manifest as a shift in soil microbial community structure and the magnitude of soil microbial activity that persists after cover crop termination, during the growth, and after the harvest of the following cash crop and such persistence can be species/cultivar specific (Cazzaniga *et al*., 2023). We also found species-specific effects on cover crop-induced soil microbial legacy effects. Soils previously planted with oil radish or sunflower resulted in significantly greater cumulative microbial substrate induced respiration, compared to soil planted with buckwheat or clover (Figure 2). Cazzaniga et al., (2023) assessed the legacy effects of cover crops on soil microbial footprints and found that, while effects of some other species persist only until the onset of cash crop, oil radish impacted soil microbial footprints until the harvesting of the cash crop. Specifically, they found that oil radish boosted both the presence and activity of microbial groups known for supressing soil-borne fungal diseases of plant. In our experiment, the cover crop species did not significantly affect abundance of total bacteria (16S rRNA gene) or total fungi (ITS region gene), but we did find that plots previously planted with radish had the highest abundance of the bacterial β-glucosidase gene (GH3) (Figure S-9) and the lowest relative abundance of the fungal β-glucosidase gene (GH3) (Figure S-11), implying evidence of negative impacts of radish on fungal communities involved in the hydrolysis of cellobiose, as previously observed when growing brassicas (Vukicevich et al., 2016; Tagele et al., 2021). We also observed greater concentrations of total N in plots previously planted with clover, which could be evidence of the legacy of N fixation by leguminous cover crops (Crotty and Stoate, 2019;Castellano-Hinojosa and Strauss, 2020).

### The influence of drying and rewetting cycles on substrate induced respiration

We found that substrate induced respiration increased after multiple drying and rewetting cycles (Figure 1). This finding is in contrast to our hypothesis that drying and rewetting cycles would supress the substrate induced respiration. Soil respiration after rewetting is often explained more by substrate supply than by the soil microbial biomass (Wang *et al*., 2003). When soils face drought conditions the soil microorganisms produce extracellular polymeric substances and osmolytes; low molecular weight organic compounds that help them maintain cell integrity (Wood et al., 2001; Kakumanu et al., 2019; Bogati & Walczak, 2022). Upon rewetting, the osmolytes, particularly trehalose, can be rapidly metabolised, either by the organisms that synthesised them or other microorganisms emerging from dormancy (Warren, 2014;Warren, 2016;Warren, 2020), and this may result in priming of soil organic matter (Allison, 2023). The role of osmolytes in drought tolerance of the soil microbial community or their function may help explain the results we observed. The greater the number of drying and rewetting cycles, the greater the substrate induced respiration. We postulate that multiple drying and rewetting cycles results in microorganisms becoming more adapted to drought (Preece et al., 2019; Evans and Wallenstein, 2012). This adaptation could be due to selection for microorganisms more capable of undergoing dormancy or synthesising osmolytes and extracellular polymeric substances as a form of ecological memory due to genetic (Canarini *et al*., 2021) or non-genetic (Riber & Hansen, 2021) diversity that increases the ability for individuals to enter and emerge from dormancy over time in populations exposed to repeated rounds of drying and rewetting cycles. Microorganisms may also be able to re-acquire osmolytes synthesised in response to the previous drying event or acquire compatible compounds already present in the soil solution (Malik & Bouskill, 2022). This greater labile organic resource would have resulted in more carbon available to respire upon wetting and a greater potential to prime the decomposition of the barley shoot powder.

### The legacy of cover crops on the adaptation of soil microbial function to repeated drying and rewetting cycles

We observed significantly lower %C in plots which had previously been planted with cover crops than control plots where cover crops were not established (Tabe 1). This finding contrasts with the results of a global meta-analysis which found that including cover crops within agricultural rotations increased %C, depending on soil texture, and the increased %C positively correlated with levels of mineralisable carbon and nitrogen in soil (Jian *et al*., 2020). However, we observed no significant difference in the concentration of CWEOC in cover cropping plots (227.6 µg C g^-1^ soil), compared to control plots (224.5 µg C g^-1^ soil), despite differences in total %C. It is therefore likely that a much greater proportion of the %C in the soils where cover crops were grown is labile. It is expected that if a drying and rewetting event is preceded by greater availability of labile C then the microorganisms have more resource available to invest in the production of osmolytes and extracellular polymeric substances to help protect them from desiccation (Kakumanu *et al*., 2019). This means that there is more carbon available to respire upon wetting and prime the decomposition of the barley shoot powder.

### The legacy of warming on the adaptation of soil microbial function to repeated drying and rewetting cycles

Soil warming is known to result in faster metabolism of microbially available organic carbon (Bradford et al., 2008; Adekanmbi et al., 2022). The observed decrease in substrate induced respiration after 8 dry/wet cycles relative to 0 dry wet cycle due to warming may be due to warming-induced depletion of dissolved organic carbon as a legacy effect. Our finding therefore implies that there will be lower soil microbial activity when soil previously exposed to warming is further exposed to extremes of drying and rewetting events, whereas cover crops may help sustain soil microbial activity after drying and rewetting. In addition, the legacy effect of warming resulted in an increase in the abundance of genes encoding Fungal β-glucosidase (GH1 gene copies g^-1^). This result could mean that, warming depleted substrate availability and stimulated the fungal community to produce extracellular enzymes, as observed by (Hou *et al*., 2016).

## Conclusions

We found that cover crop mixtures created a negative legacy effect in the soil which resulted in lower cumulative substrate induced respiration than expected from the average of the same species grown in monoculture. However, some monocultures (radish and sunflower) left a greater positive legacy on substrate induced respiration than others (buckwheat and clover). Exposure of the soil to multiple drying and rewetting cycles resulted in greater substrate induced respiration and this response was increased by the legacy of cover crops and decreased by the legacy of soil warming. It is likely that exposure of soils to drying and rewetting cycles induces the microbial synthesis of osmolytes and extracellular polymeric substances to help microorganisms avoid desiccation during periods of dormancy and selects for a microbial community more capable of entering and emerging from dormancy. There was a positive correlation between the cumulative respiration after 8 wet/dry cycles relative to cumulative respiration after 0 wet/dry cycles and CWEOC, highlighting the importance of substrate availability for microbial adaptation to drying and rewetting cycles. Labile carbon in soils previously planted with cover crops may provide the substrates to synthesise osmolytes and extracellular polymeric substances, whereas the depletion of labile carbon by warming decreases the substrates available to synthesise these compounds. Therefore, cover crops may be able to mitigate the impact of warming on the ability for soil microorganisms to adapt to drying and rewetting cycles.

## Authorship

Adetunji Alex Adekanmbi – Conceptualization, Data curation, Formal analysis, Investigation, Methodology, Visualisation, Writing – original draft

Yiran Zou – Investigation, Methodology, Writing – review & editing Xin, Shu - Investigation, Methodology, Writing – review & editing

Giacomo, Pietramellara - Funding acquisition, Resources, Supervision, Writing – review & editing

Shamina Imran, Pathan - Data curation, Formal analysis, Investigation, Methodology, Writing – review & editing

Lindsay, Todman - Conceptualization, Methodology, Formal analysis, Writing – review & editing

Tom Sizmur - Conceptualization, Funding acquisition, Project administration, Supervision, Visualisation, Writing – original draft

## Acknowledgements

The authors acknowledge the technical support provided by Richard Casebow, Caroline Handley, Mike Charij, Francesco Leva, Anne Dudley, Fengjuan Xiao, Marta O’Brian, Laurence Dale, Caroline Thomas, Luci Corbett

