## Supplementary Information for "Legacy of warming and cover crops on the response of soil microbial function to repeated drying and rewetting cycles"

This Supporting Information file contains 11 Figures and 1 Table

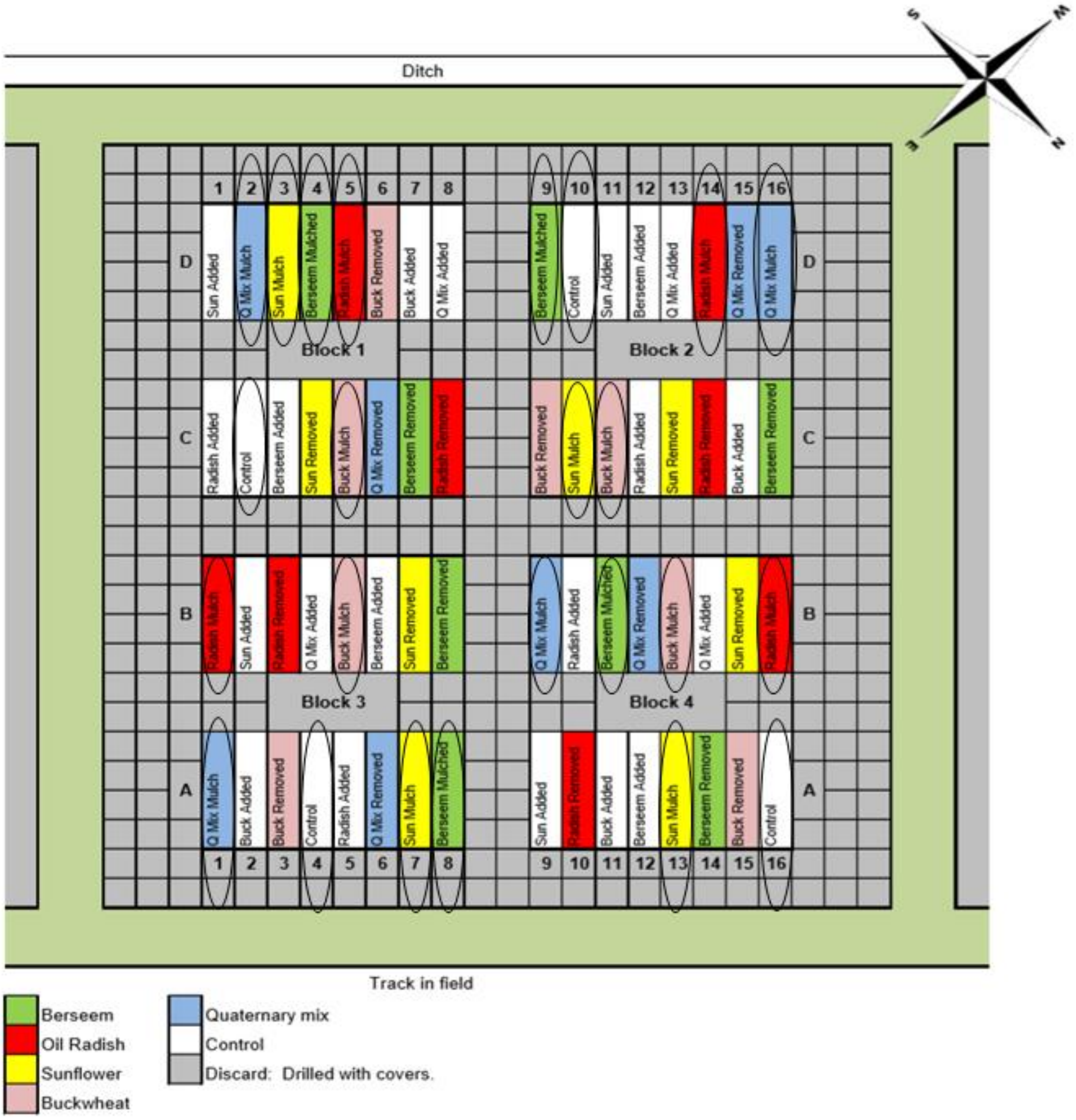


**Figure S-1: Experimental layout of the field experiment located at Sonning Farm, in Berkshire, UK. Circled plots indicate those upon which Open Top Warming Chambers were placed on the southwest end of the plot. On each of these 24 plots cover crops (as a single species monoculture, or a 4-species mixture) were grown and then mulched and the residues incorporated into the soil prior to sowing the next cash crop. On other plots (not used in this study) residues were removed and added to other plots prior to incorporation.**

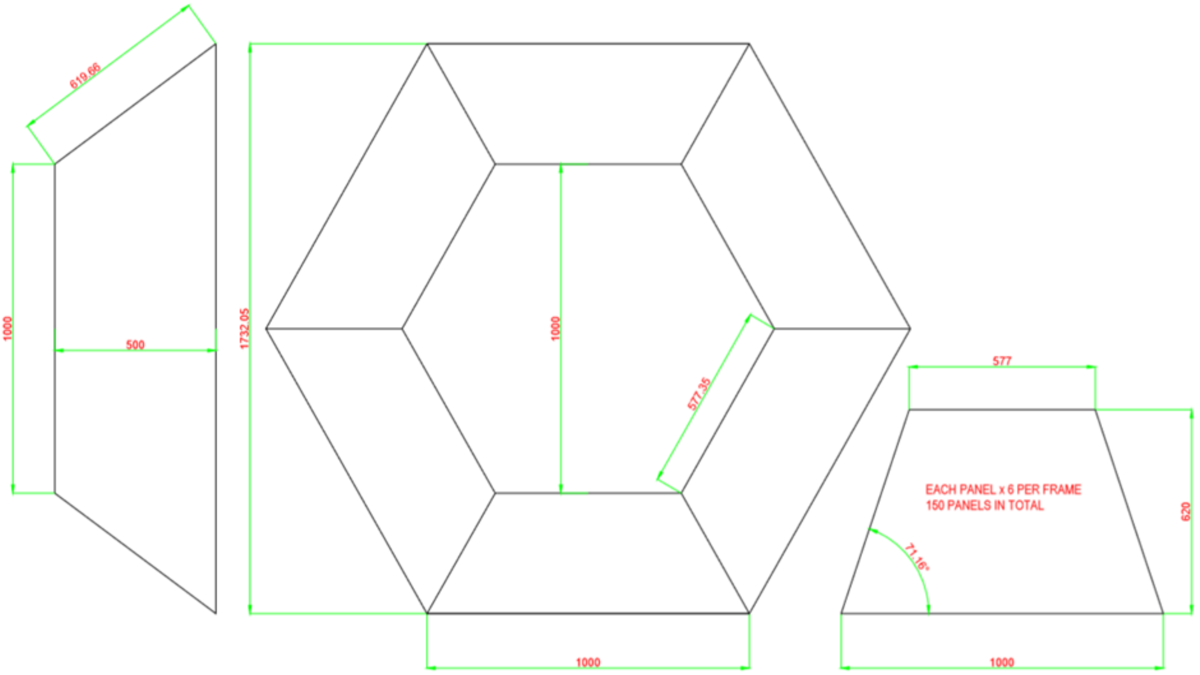


**Figure S-2: Schematics of the Open Top Warming Chambers deployed on plots. Units of dimensions are mm. Credit: Mike Charij**

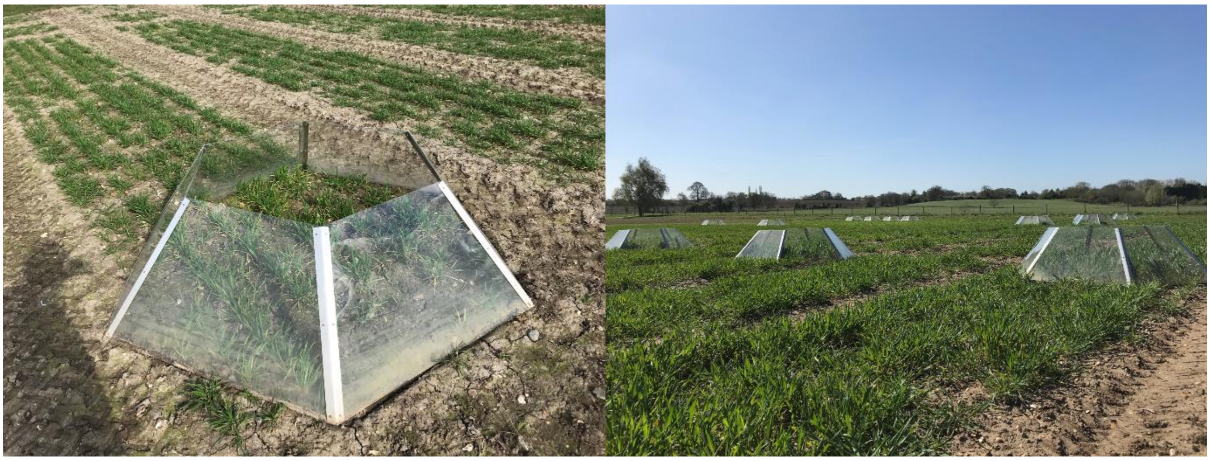

**Figure S-3: Images showing the installed chambers**

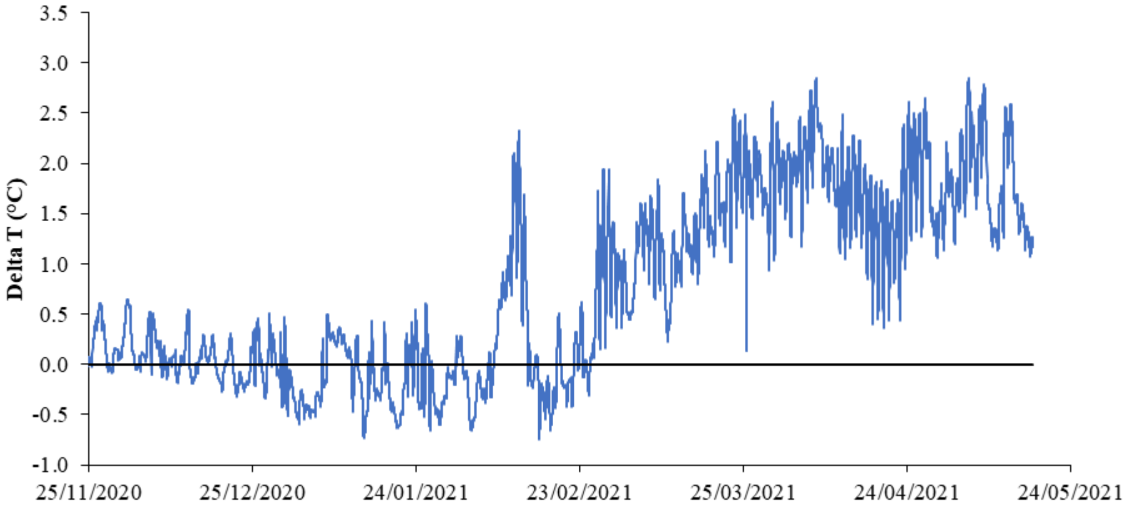


**Figure S-4: The difference in temperature recorded in soil underneath an open top passive warming chamber and the equivalent location at the other end of the same plot. The horizontal line represents no warming (Delta T = 0 °C). Positive data indicates warming caused by the chamber and negative data indicates cooling caused by the chamber, relative to the ambient soil temperature**

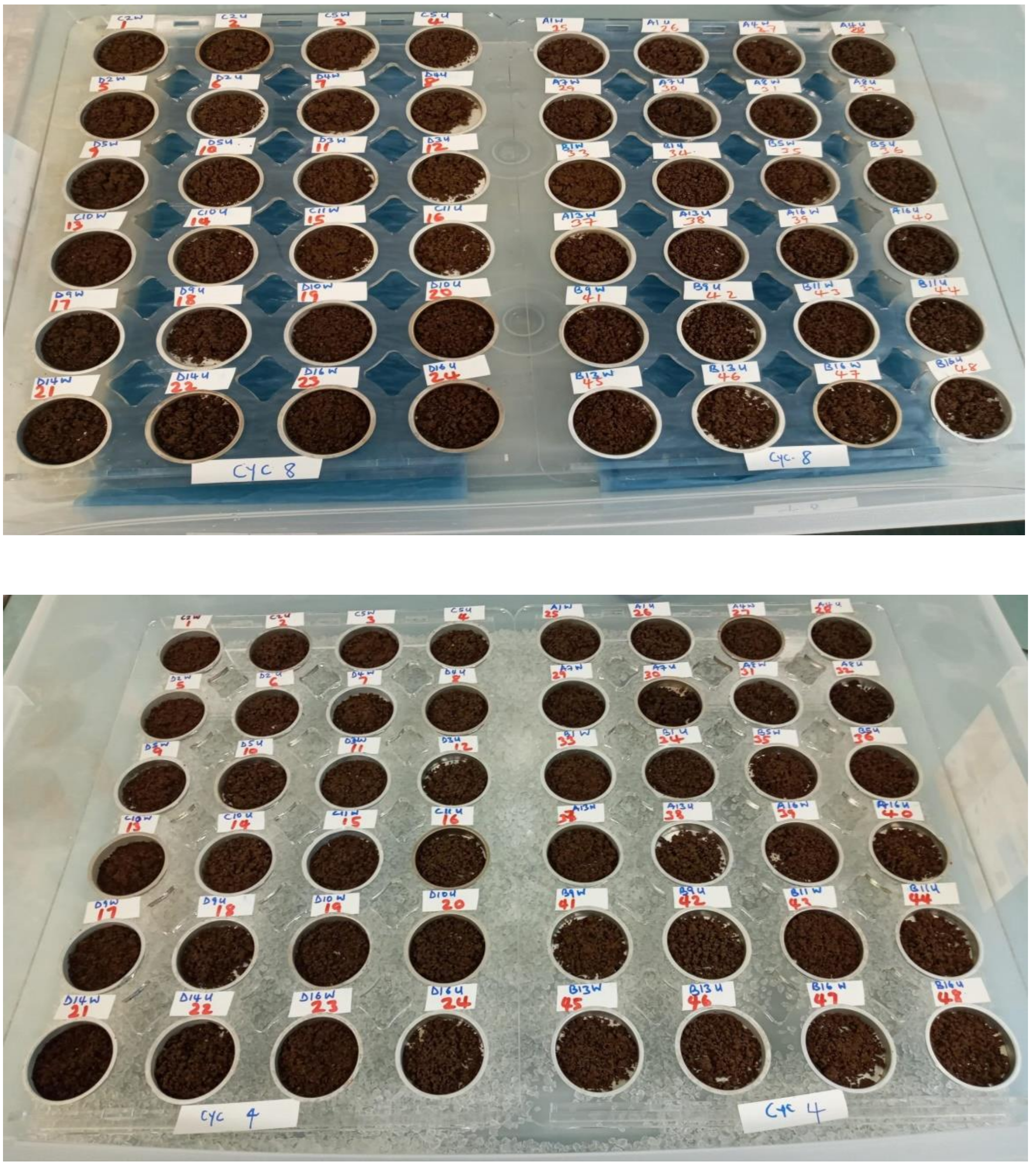


**Figure S-5: Images showing the placement of soil samples above wet paper towel (top panel) and desiccant (bottom panel) during the imposition of wet/dry cycles.**

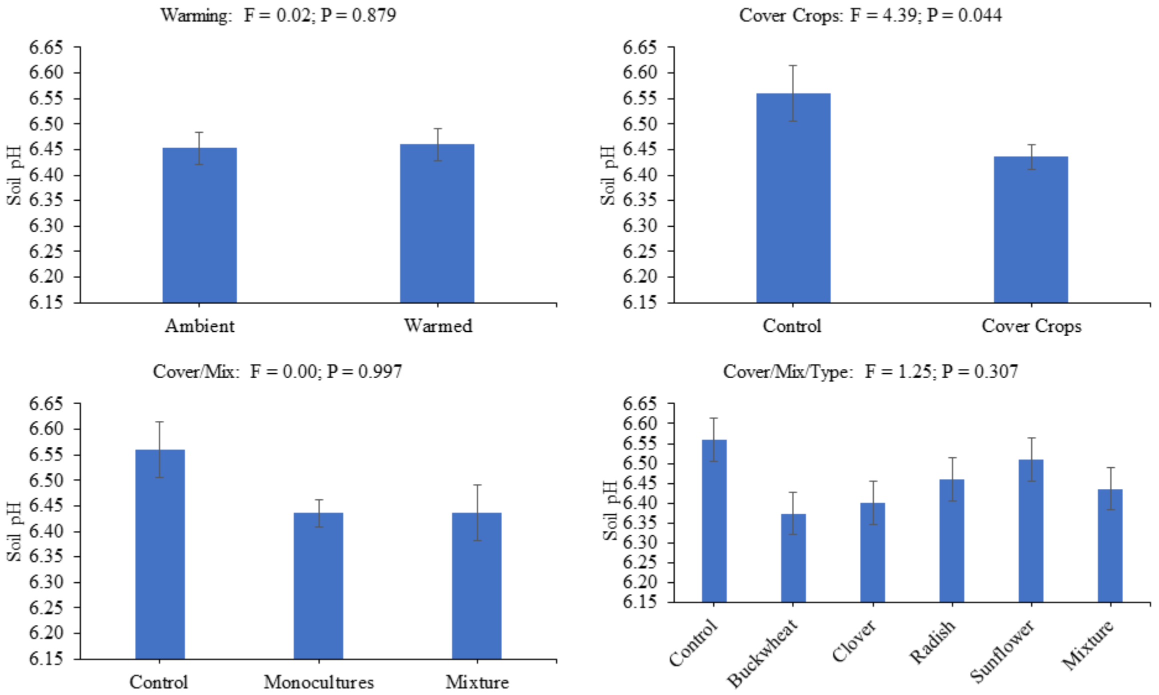


**Figure S-6 : Effects of experimental factors (Warming, Cover, Mix, and Type) on Soil pH**

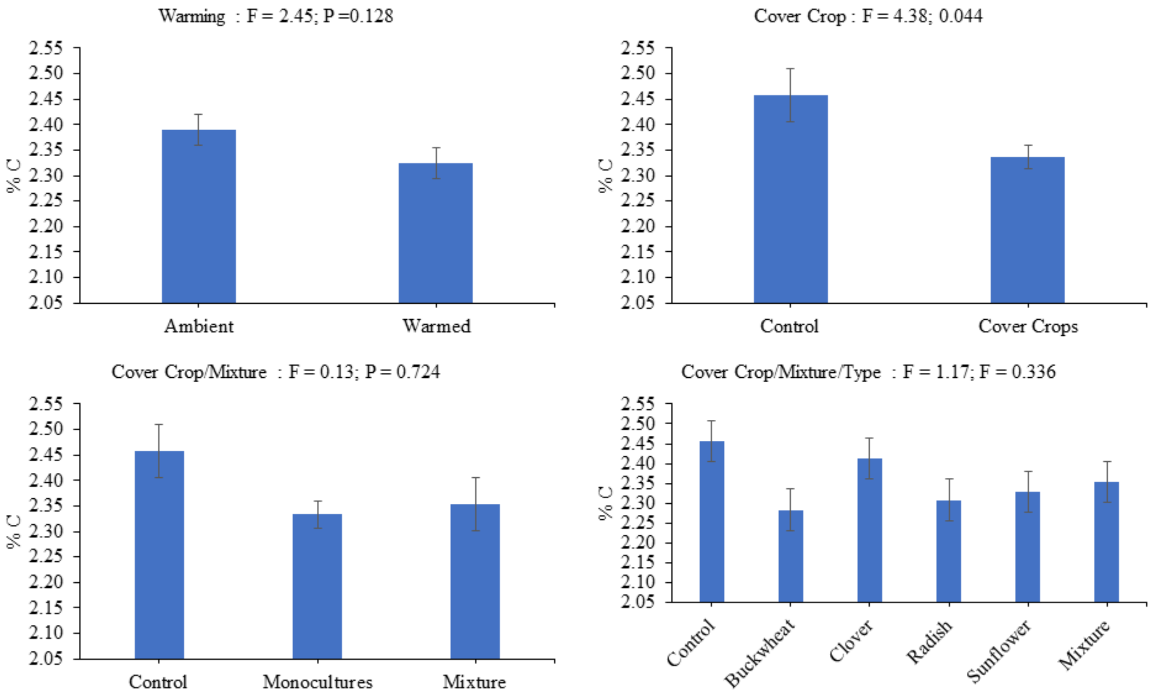


**Figure S-7: Effects of experimental factors (Warming, Cover, Mix, and Type) on Total Carbon (%C)**

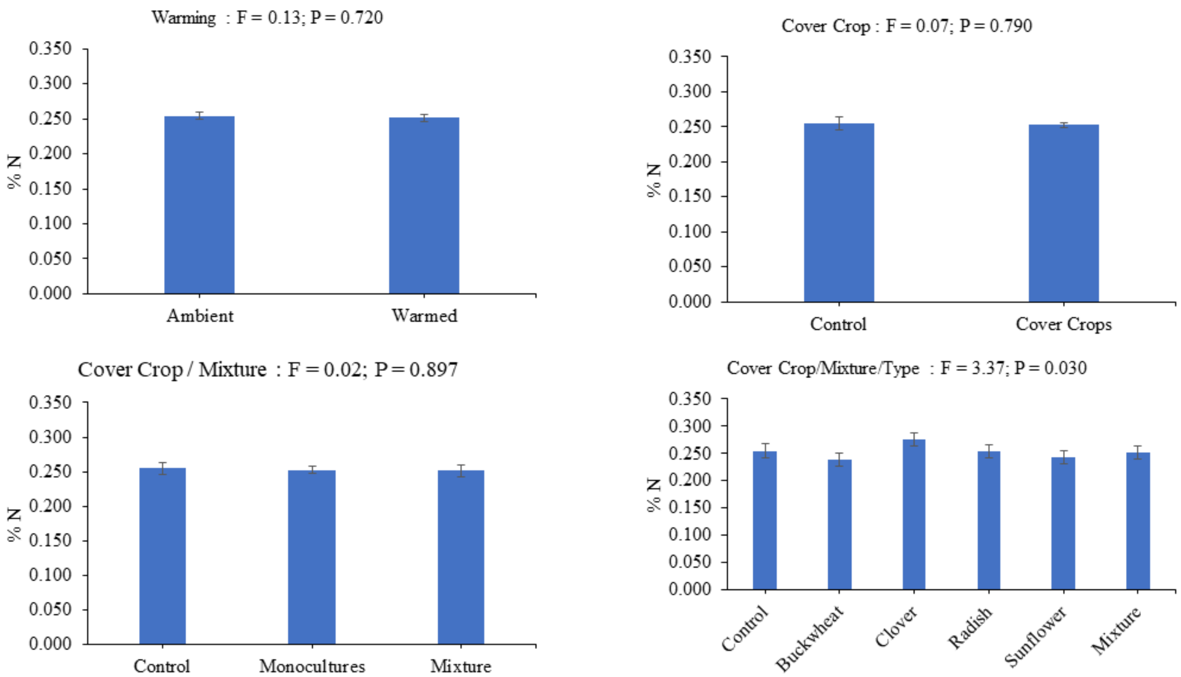


**Figure S-8: Effects of experimental factors (Warming, Cover, Mix, and Type) on Total Nitrogen (% N)**

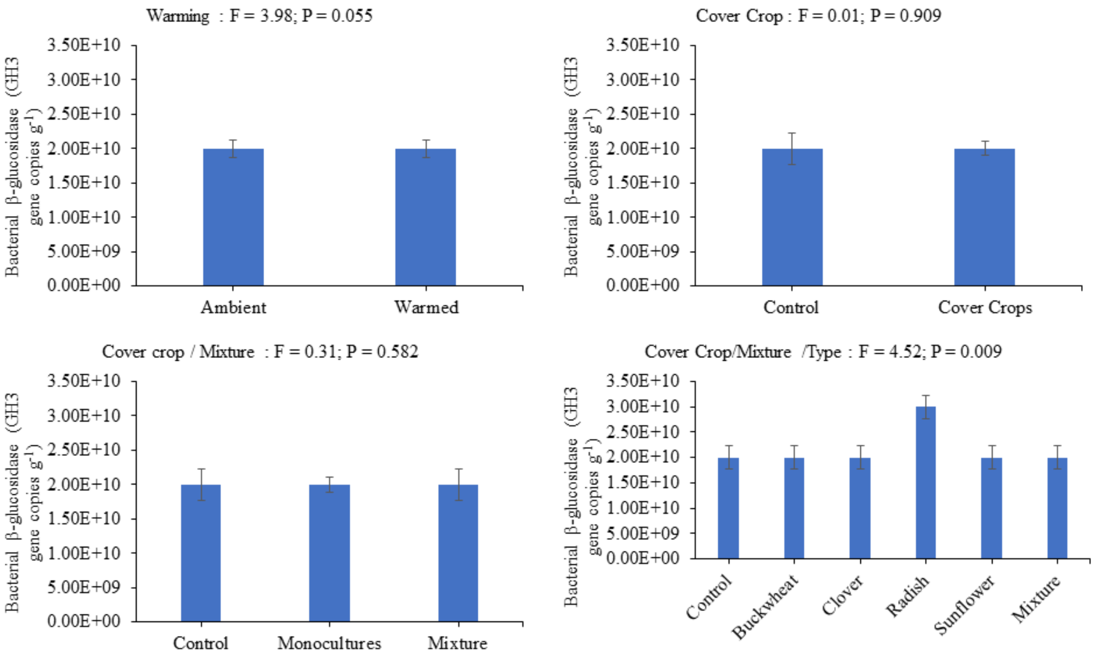


**Figure S-9: Effects of experimental factors (Warming, Cover, Mix, and Type) on relative abundance of gene encoding Bacterial β-glucosidase (GH3 gene copies $g^{-1}$)**

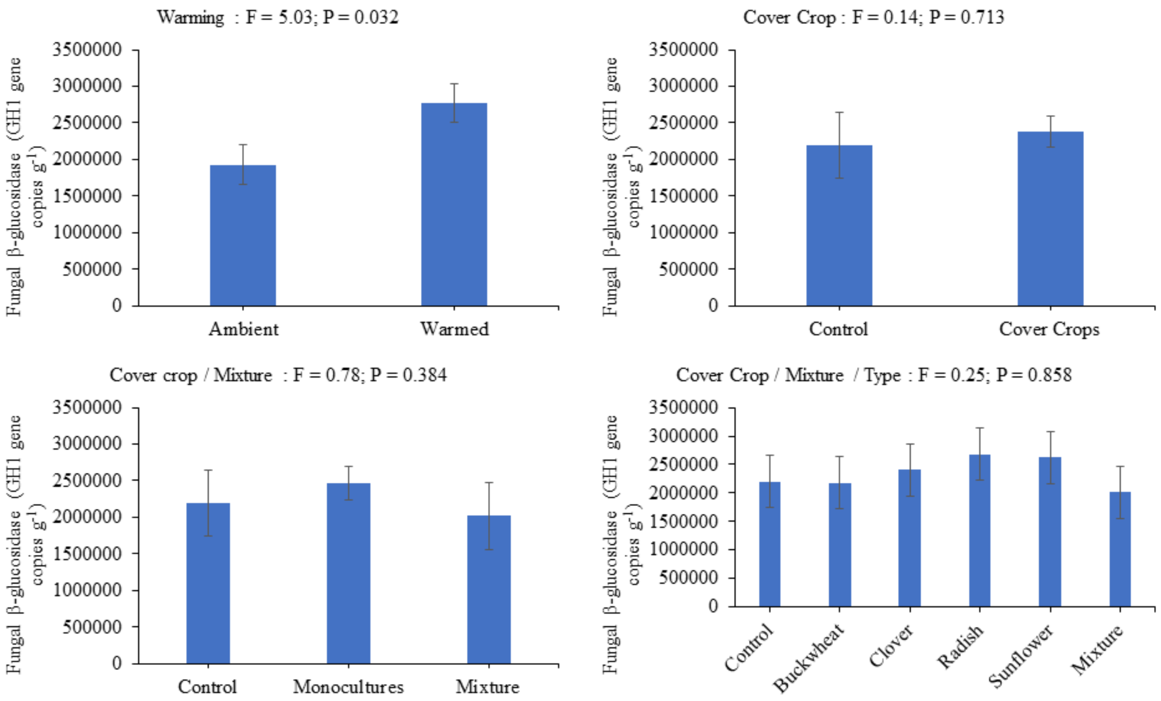


**Figure S-10: Effects of experimental factors (Warming, Cover, Mix, and Type) on relative abundance of gene encoding Fungal β-glucosidase (GH1 gene copies $g^{-1}$)**

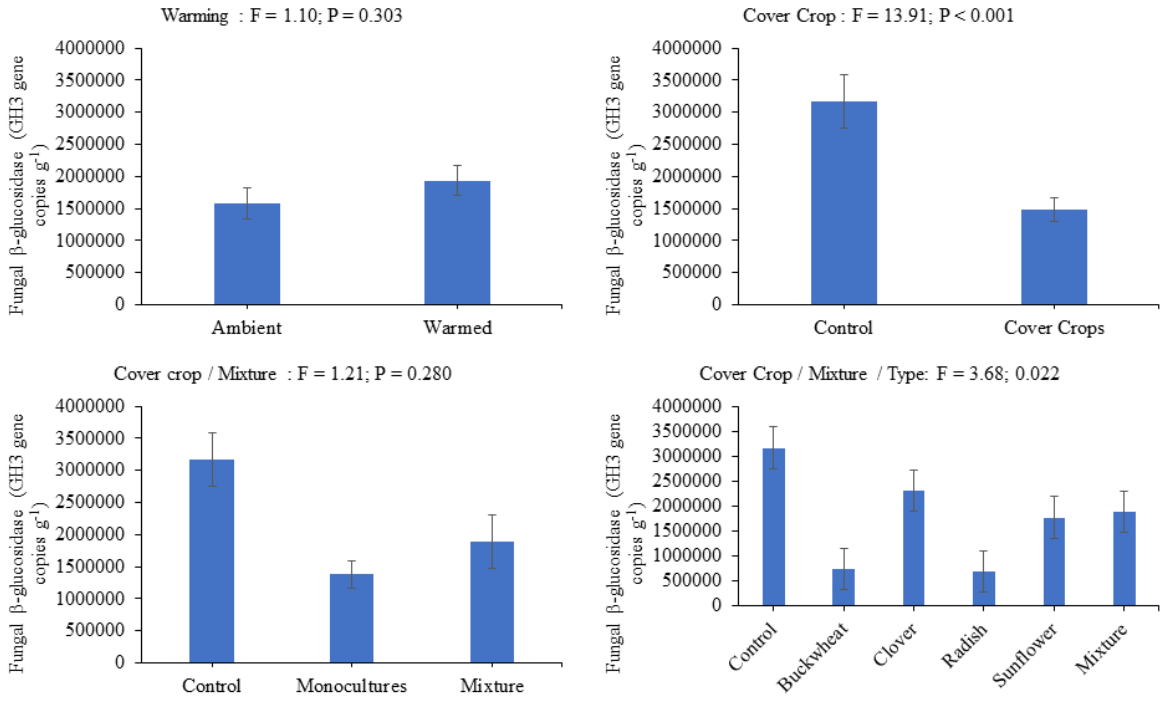


**Figure S-11: Effects of experimental factors (Warming, Cover, Mix, and Type) on relative abundance of gene encoding Fungal β-glucosidase (GH3 gene copies $g^{-1}$)**

**Table S-1 Quantitative PCR primers used to quantify phylogenetic and functional microbial diversities in this study.**

| Target Groups | Gene | Primer Name/Sequence (5'-3') | Amplicon Length (bp) | Reference |
|---|---|---|---|---|
| **Bacteria** | 16S rRNA | EUB341F: CCTACGGGAGGCAGCAG<br>EUB515R: TACCGCGGCKGCTGGCA | 174 | (Muyzer *et al.*, 1993) (Simmons *et al.*, 2007) |
| **Fungi** | ITS2 | ITS3: GCATCGATGAAGAACGCAGC<br>ITS4: TCCTCCGCTTATTGATATGC | 386 | (White *et al.*, 1990) |
| **GH1-Bacteria** (β-glucosidase) | β-glucosidase | β-gluF2: TTCYTBGGYRTCAACTACTA<br>β-gluR4: CCGTTYTCGGTBAYSWAGA | 180 | (Cañizares *et al.*, 2011) |
| **GH3-Bacteria** (β-glucosidase) | β-glucosidase | BGH3BF: TTCGGCGAAGAYCC<br>BGH3BR: ACGCCTTYRWARCC | 300 | (Li *et al.*, 2013) |
| **GH1-Fungi** (β-glucosidase) | β-glucosidase | bglFGH1F: TGGATCNTTCAAYGARCC<br>bglFGH1R: GTAGTGGTTCAGCCRWARAA | 350 | (Pathan *et al.*, 2017) |
| **GH3-Fungi** (β-glucosidase) | β-glucosidase | bglFGH3F: GTTCCGTCATGTGCTCYTAYAA<br>bglFGH3R: CATGATACGGGTAGCCATRTC | 300 | (Pathan *et al.*, 2017) |
